# Bayesian hierarchical models can infer interpretable predictions of leaf area index from heterogeneous datasets

**DOI:** 10.1101/2021.09.20.461084

**Authors:** Olivera Stojanović, Bastian Siegmann, Thomas Jarmer, Gordon Pipa, Johannes Leugering

## Abstract

Environmental scientists often have to predict a complex phenomenon from a heterogeneous collection of datasets. This is particularly challenging if there are systematic differences between them, as is often the case. Accounting for these differences requires a larger number of parameters and thus increases the risk of overfitting. We investigate how Bayesian hierarchical models can help mitigate this problem by allowing the practitioner to explicitly incorporate information about the dataset structure and general domain knowledge. To this end, we look at a typical application in remote sensing: the estimation of leaf area index (of white winter wheat), an important indicator for agronomical modeling, from measurements of reflectance spectra collected at different locations and growth stages. Since the insights gained from such a model could be used to inform policy or business decisions, the interpretability of the model is a primary concern. We, therefore, focus on models that capture the association between leaf area index and the spectral reflectance at various wavelengths by spline-based kernel functions, which can be visually inspected and analyzed. We compare models with three different levels of hierarchy: a non-hierarchical baseline model, a model with hierarchical bias parameter, and a model in which bias and kernel parameters are hierarchically structured. We analyze them using Markov Chain Monte Carlo sampling diagnostics and an intervention-based measure of feature importance. The improved robustness and interpretability of this approach lead us to recommend Bayesian hierarchical models as a versatile tool for environmental sciences and beyond, particularly in scenarios where the available data sources are heterogeneous.

## Introduction

The leaf area index (LAI) is a unitless measure (m^2^ m^−2^) of the one-sided leaf surface area of a plant relative to the soil surface area. [1] It characterizes, among other variables, photosynthetic performance of plants [1, 2, 3], and the size and density of the crop’s canopy and thus serves as an important indicator for the plant’s growth stage and stand productivity [4, 5, 6, 7, 8]. Therefore, it plays a major role in meteorological, ecological, and agronomical modeling [9, 10, 11, 12, 13, 14], as well as for studying the influence of climate change on crop growth. [15, 16]

Various non-destructive methods exist to measure or estimate LAI directly [17], but they typically require taking a large number of manual measurements in the field. Since this is a laborious process and it can be difficult to control for confounding variables such as weather, alternative faster approaches to infer LAI from indirect measures, e.g., spectroscopy, have been investigated [5]. Spectral reflectance curves have been well understood and are relatively easy to measure. Since they can also be acquired through aerial or satellite surveillance, this could greatly simplify monitoring crop growth across large or remote areas [18].

However, the spectral reflectance is also affected by countless other factors, such as the crop type, phenology, sun illumination, local micro-climate, the type of soil, or the amount of precipitation [19]. The relationship between LAI and reflectance spectrum may also vary throughout the life-cycle of the crop. A model that predicts LAI from reflectance spectra should be adjusted to these specifics, but this poses a practical challenge: the potential confounding variables are too numerous to include in the model, and instead, we have to account for them implicitly by allowing the model to vary across different datasets. In addition, the amount of labeled training data that can be acquired for each field and/or season is limited, so it is inefficient, if not impossible, to fit a separate model for each field and/or growth stage. It is possible, in principle, to train a single model that generalizes across these conditions by simply pooling multiple datasets that were acquired under different conditions (see [20], for example). However, since all of the specifics of the individual dataset are lost in the process, such a model is likely to perform worse than a field- and growth-stage-specific model could, given sufficient training data. Ideally, we would like to find a compromise between these two extremes that allows us to generalize over all the available datasets yet makes specifically adjusted predictions for each dataset.

We present a solution to this problem in the form of a hierarchical, parameter-efficient Bayesian model. Our model learns an easily interpretable general relationship between reflectance spectra and LAI, as well as the dataset-specific deviations from that baseline. We identify relevant spectral regions and quantify their contribution to the prediction of LAI. By using a variant of Markov-Chain-Monte-Carlo (MCMC) sampling, we can incorporate domain knowledge or regularization through prior distributions of the parameters and provide a full posterior probability distribution over these parameters, which allows the quantification of uncertainty. We compare two variants of this hierarchical approach with a non-hierarchical alternative and find that it indeed offers a favorable trade-off between prediction accuracy and model complexity.

Bayesian hierarchical models similar to the one suggested here are especially appealing for environmental sciences [21], where they have seen increasingly widespread use. For example, several recent studies applied Bayesian hierarchical models to time series of multispectral satellite images in order to assess the effects of climate change through land surface phenology [22, 23, 24], or other indicators such as Normalized Difference Vegetation Index [25]. A similar remote sensing approach has been used to predict LAI and its spatio-temporal evolution for bamboo [26] and other forests [27, 28]. For agronomical models of the LAI of food crops such as rice [29], Brazilian Cowpea [30] or white winter wheat [31], local multispectral measurements are often used instead of – or in addition to – satellite images. In most of these studies, Bayesian hierarchical models are used to impose prior domain knowledge, combine multi-model data sources, and integrate data collected at multiple resolutions of space and/or time, all of which ultimately improve prediction performance. By contrast, our primary goal is to show how Bayesian hierarchical models and associated tools can be used from a data science perspective to construct and diagnose simple and interpretable models, which can cope with the kind of heterogeneous datasets often encountered in environmental sciences.

## Methods

### The dataset

We evaluate our proposed model on a combination of four datasets, totalling 191 pairs of measured reflectance spectra (see also S1 Fig for examples) and corresponding measurements of the LAI on fields of white winter wheat (lat. *Triticum aestivum*). The recorded spectra cover wavelengths in the range from 350 nm to 2500 nm, of which we use the range from 400 nm to 1350 nm (for details see 1). We preprocess these spectra by smoothing with a first-order Gaussian filter with width *σ* = 10 nm. Each pair of measurements was taken on a different square plot of size 50 cm × 50 cm. The LAI values of each plot were measured multiple times in a non-destructive way and averaged to a single value per plot. Five reflectance spectra were acquired and averaged for each plot using a spectroradiometer from a height of 1.4 m above ground with a nadir view and converted to absolute reflectance values using a reflectance standard of known reflectivity. The data were collected on four different fields in different years, corresponding also to different stages in the plants’ growth cycle, and there are minor differences in the data collection procedure.

The first two sampling areas, which we call Field A and Field B in the following, are located near Köthen, Germany, which is a part of one of the most important agricultural regions in Germany. The region is distinctly dry, with 430 mm mean annual precipitation due to its location in the Harz mountains. The mean annual temperature varies between 8 °C to 9 °C. The study area has an altitude of 70 m above sea level and is characterized by a Loess layer up to 1.2 m deep that covers a slightly undulated tertiary plain. The predominant soil types of the region are Chernozems, in conjunction with Cambisols and Luvisols. At two locations in this region, 57 measurements were recorded on 7th to 8th May 2011, and another 74 on 24th to 25th May 2012. In 2011 and 2012, respectively, 25° and 14° field of view optics were used for the spectroradiometer, and five and four LAI measurements were averaged per plot. Data from this study area was also used and described in more detail in [20].

The other two sampling areas, called Field C and Field D in the following, are located near Demmin, Germany, with a mean annual precipitation of 550 mm, and a mean annual temperature of 8 °C. Albeluvisols interspersed by Haplic Luvisols dominate the sand-rich area. The observed area is south of the river Tollense, where the ground elevation drops from 70 m to 7 m due to glacial moraines causing high variability in soil conditions. At two locations in this region, 26 measurements were recorded on 5th June 2015, and another 34 on 10th May 2016. In this case, six measurements of LAI were averaged for each plot.

For a summary of these parameters, see table 1.

**Table 1.** Parameters of the dataset and the collection procedures.

|  | Field A | Field B | Field C | Field D |
| --- | --- | --- | --- | --- |
| collection date | 7th to 8th May 2011 | 24th to 25th May 2012 | 5th June 2015 | 10th May 2016 |
| measurements | 57 | 74 | 26 | 34 |
| location | Köthen, Germany |  | Demmin, Germany |  |
| LAI device | SS1 SunScan | LAI-2000 | LAI-2000 |  |
| spectral device | ASD FieldSpec III | SVC HR-1024 | SVC HR-1024i |  |
| field of view | 25° | 14° | 14° |  |
| measurement height | 1.4 m above ground |  |  |  |
| reflectance standard | Spectralon, Labsphere Inc., USA |  |  |  |
| spectral range measured | 350 nm to 2500 nm |  |  |  |
| spectral range used | 400 nm to 1350 nm at a resolution of 1 nm, smoothing with $\sigma = 10$ nm | | | |
| LAI range | 0.5 to 3.32 | 1.14 to 6.16 | 1.72 to 7.46 | 0.48 to 5.25 |

In the following, we will reference specific subsets of this data; hence we introduce some notation here: We assume that all spectra-LAI-pairs are numerically indexed in the range *J* = {1, …, 19}, and we defined four index sets *J*^1^, *J*^2^, *J*^3^, *J*^4^ ⊂ *J* that correspond to the measurements from Field A, Field B, Field C, and Field D, respectively. We denote the *i*^th^ reflectance spectrum from dataset *j* by the function 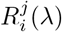 of the wavelength *λ* ∈ [400 nm, 1350 nm], and the corresponding measured LAI value by 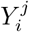.

### Feature extraction from reflectance spectra with a spline basis

The data collection and preprocessing steps outlined above result in reflectance spectra of wavelengths 400 nm to 1350 nm at a resolution of 1 nm. Since this representation is much too high dimensional for direct use, we extract the most important information into a low dimensional representation by computing the inner product between the preprocessed reflectance spectra and a set of eleven cubic basis splines (B-splines) with adaptively placed knots (see figure 1). The positions *κ*_*i*_, *i* ∈ {1, …, 10} of the knots, which determine the shape of the individual basis splines, are chosen such that the cumulative absolute curvature *Q*(*κ*_*k*+1_) − *Q*(*κ*_*k*_) of the average reflectance spectrum is equal between any two successive knots *k* and *k* + 1. We compute the absolute curvature *q*(*λ*) by convolving^1^ the average reflectance spectrum 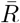 with the second derivative of the Gaussian function *g*, and then compute the absolute value thereof. Formally, we can express this as follows:

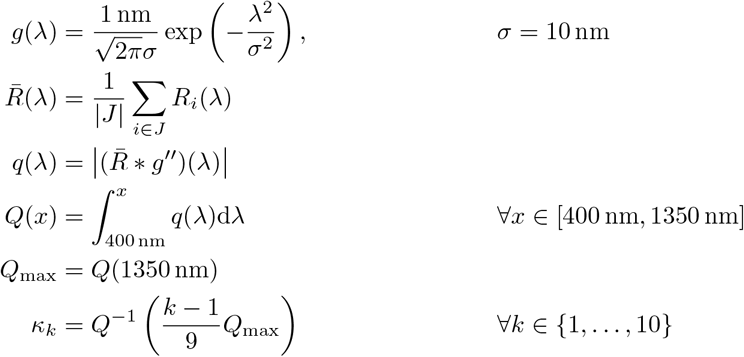

**Figure 1.**
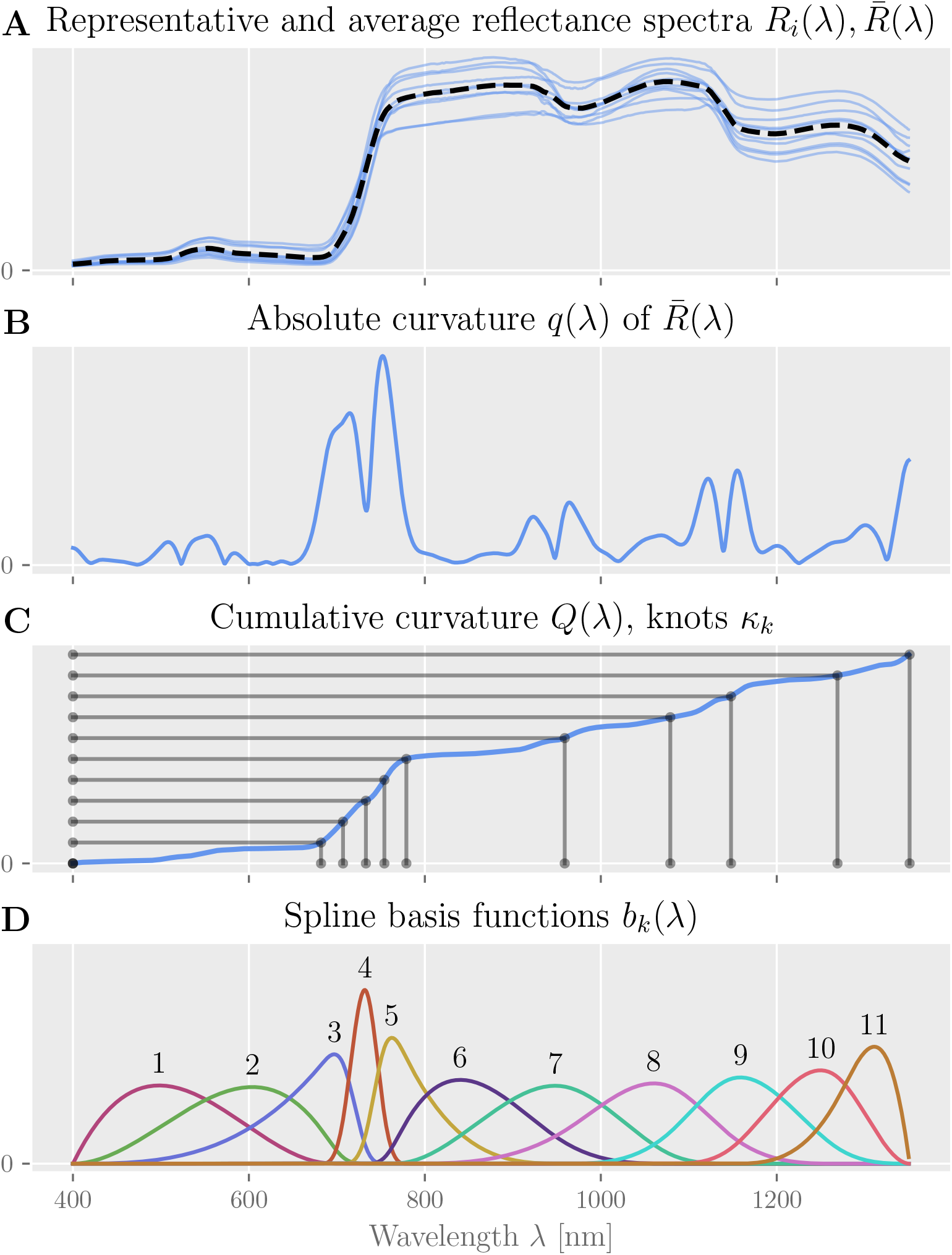
Adaptive knot-placement for B-Splines. (**A**) For the measured reflectance spectra *R*_*i*_(*λ*), (**B**) we calculate the mean absolute curvature *q*(*λ*). (**C**) We then find knot positions such that the integral *Q*(*λ*) of this measure between any two successive knots *κ*_*k*_, *κ*_*k*+1_ is identical. (**D**) The result are 11 cubic spline basis functions *b*_*k*_(*λ*) with non-uniformly spaced knots

The eleven basis functions *b*_*k*_ are generated from this knot-vector *κ* using the standard Cox-DeBoor algorithm [32], where the multiplicity of the first and last knot is three, i.e., all basis functions go to zero at their respective start and end knots.

This heuristic algorithm results in a proportionally larger number of knots, and thus higher spatial resolution, where the reflectance spectra have the largest absolute curvature and hence “have most structure”; see also [33, 34, 35]. For each of the data-subsets *J*^*j*^, *j* ∈ {1, …, 4}, we can then compute our model’s *feature* or *design matrix X*^*j*^ using these basis functions *b*_*k*_(*λ*):

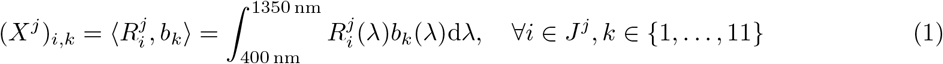

### Bayesian Markov-Chain-Monte-Carlo regression for predicting LAI

Our primary objective is to construct a simple, interpretable model that can reliably predict the LAI value of a wheat plot directly from a corresponding reflectance spectrum. We are additionally interested in analyzing the model’s confidence, how well it generalizes, and which features it relies on most to make a prediction. Since the total available data is limited and stems from four heterogeneous datasets, prior constraints are required to prevent overfitting.

In order to meet all of these requirements, we design three different (non-)hierarchical Bayesian generalized linear models (GLM) [36, 37] of different complexity. For each of these, we infer a full posterior distribution over model parameters from training data and use this to provide a full posterior predictive distribution over LAI scores on testing data. To generate representative samples from these probability distributions, we use a specific type of Hamiltonian Monte Carlo sampling, namely No-U-Turn-Sampling[38] (NUTS), as implemented by the probabilistic programming package *pyMC3* [39].

#### Model 1: A baseline model with pooled data

As a baseline (see figure 2), we construct a simple generalized linear model, which we apply to all of the datasets *J*^*j*^, *j* ∈ {1, … 4} together. This model merely pools all available data but does not account for any systematic differences that might exist between the individual datasets. We assume the logarithm of the observed LAI scores to be normally distributed around an affine linear predictor *μ*^*j*^ with deviation *V*, which is a parameter of the model with log-normal prior. The predictor *μ*^*j*^ is computed by the matrix-vector product between the dataset’s feature matrix *X*^*j*^ and the model’s weight vector *w* = (*w*_1_, …, *w*_11_), plus an additional bias parameter *b*. Including the unknown deviation parameter *V*, the model thus has a total of 13 free parameters to be inferred from data. The individual parameters *w*_*k*_ and *b* have normal priors with standard deviation 1 and 11, respectively, to allow the individual bias term to counteract the effect of all 11 weights, if necessary. The baseline model is described by equation 2.

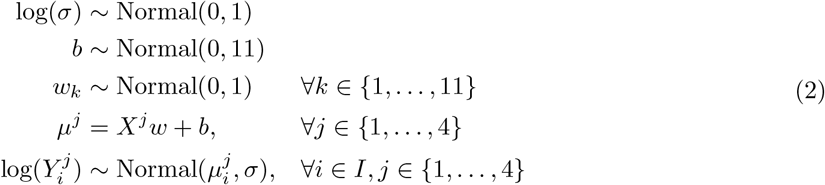

**Figure 2.**
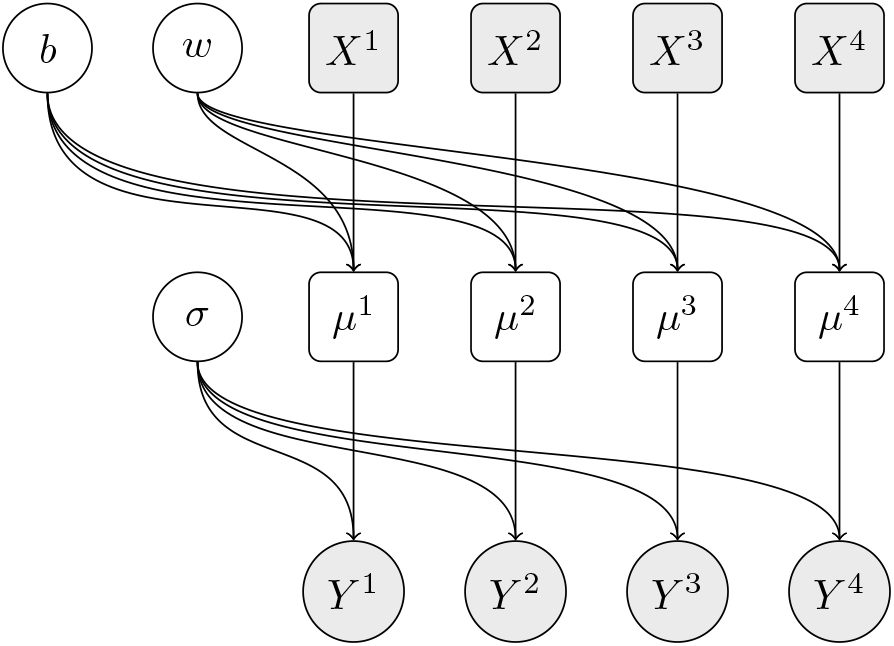
Dependency graph of the baseline model. For each dataset *j* (encoded in the feature matrix *X*^*j*^ and corresponding labels *Y* ^*j*^), the prediction depends on the same three shared parameters *b, w* and *σ*. Circles represent random variables, rectangles represent deterministic variables, filled shapes represent observed variables.

#### Model 2: A model with hierarchical bias

Our second model (see figure 3) extends the baseline model by an additional bias parameter *b*^*j*^ for each dataset, and thus has a total of 17 free parameters. Due to the logarithmic link function, this additional parameter per dataset allows accounting for the overall variation in scale between the four different datasets. But rather than setting each parameter *b*^*j*^ independently (and thus adding three full degrees of freedom), we constrain them to be clustered around a common bias value *b*^*^, which replaces the bias term *b* in the baseline model. Therefore, the prior for the new variables *b*^*j*^ is a Normal distribution centered at *b*^*^ with an order of magnitude smaller standard deviation of 11*/*10 = 1.1. The affine linear predictor *μ*^*j*^ then depends on the dataset-specific bias term *b*^*j*^. The hierarchical bias model is described by equation 3.

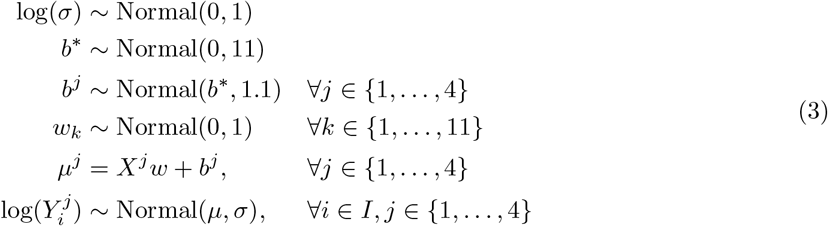

**Figure 3.**
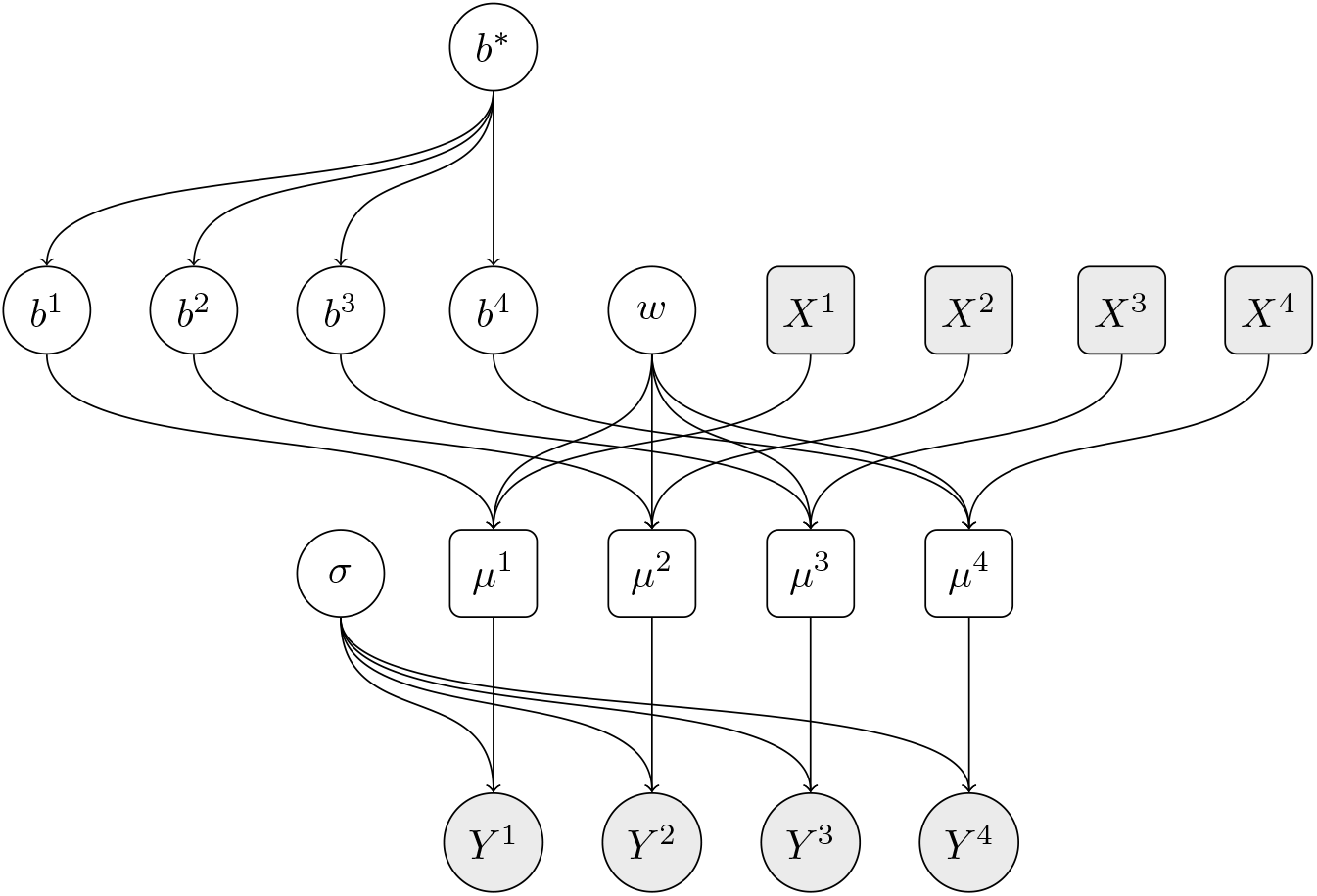
Dependency graph of the hierarchical model with dataset-specific bias terms. The predictions for each dataset *j* depend on an individual bias parameter *b*^*j*^, which in turn depends on the shared mean bias parameter *b*^*^.

#### Model 3: Full hierarchical model

Our third model (see figure 4) extends the second model even further by also allowing the model weight vector *w* to vary for each dataset. Just like we did for the bias terms, we introduce the new parameter vectors *w*^*j*^, and we constrain the individual parameters 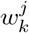 to be clustered around the corresponding common values 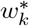 with standard deviation 0.1. This increases the model’s degrees of freedom by an additional 44 parameters (11 for each dataset), resulting in a total of 61 free parameters. The affine linear predictor *μ*^*j*^ then depends on a dataset specific weight vector *w*^*j*^ and a dataset specific bias term *b*^*j*^. The full hierarchical model is described by equation 4.

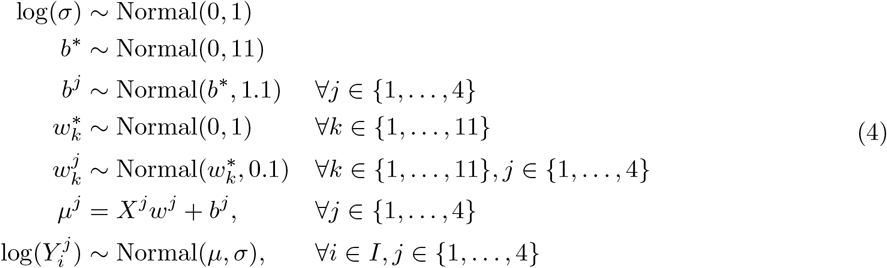

**Figure 4.**
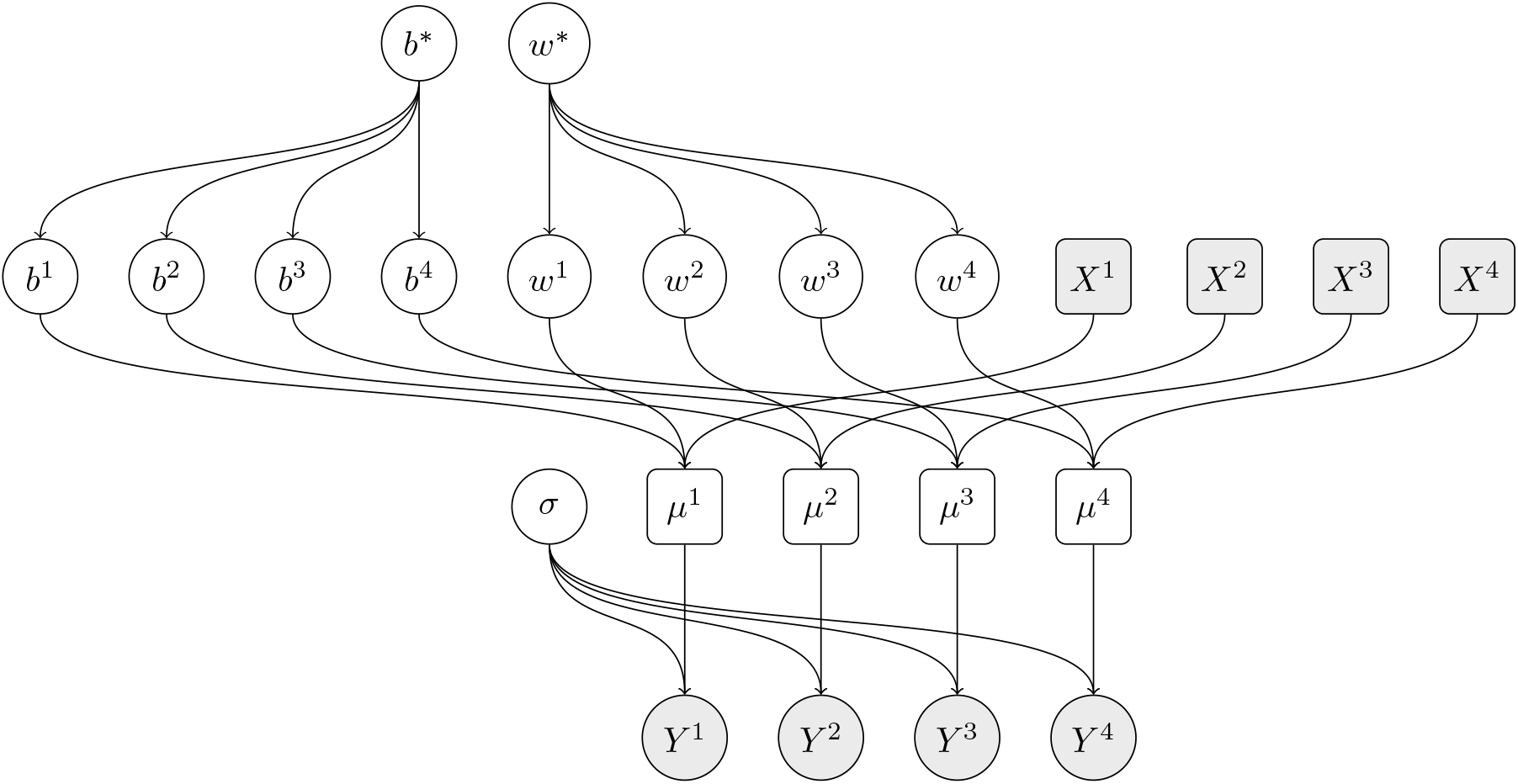
Dependency graph of the hierarchical model with dataset-specific bias and weight terms. The predictions for each dataset *j* now depend on an individual bias parameter *b*^*j*^ and weight vector *w*^*j*^, which in turn depend on the shared bias parameter *b*^*^ and the shared weight vector *w*^*^, respectively.

### Model selection using Pareto-Smoothed Importance Sampling

To get an unbiased estimate of our model’s generalization error from the very limited available data, we would like to perform leave-one-out cross-validation (LOO-CV) and compute the expected log posterior predictive density (ELPD) for new data. Unfortunately, this is a prohibitively expensive computation when combined with MCMC sampling. However, the generated samples and their associated log-likelihood values contain sufficient information to directly estimate the LOO-CV ELPD by an appropriate weighting of the samples. This procedure is called Pareto-smoothed importance sampling [40] (PSIS), and it is beneficial in situations like this, where an MCMC sampling-based model is trained on a small dataset. As a result, we get for each model the ELPD score, which we use to compare the three proposed models, a parameter *η*, which can be interpreted as the effective number of degrees of freedom in the model, and the so-called Pareto shape parameters *k*_*i*_, which assess for each data point *i* in the dataset, how much it affects the ELPD estimation. For data points where *k*_*i*_ exceeds 0.7, the PSIS-LOO-CV estimate becomes unreliable, which can also indicate an under-constrained model or an outlier in the data [40]).

### Evaluation of feature importance

To estimate the importance that our model assigns to each feature of the reflectance spectra, we calculate a model-agnostic measure of feature importance [41] called *model reliance* (MR). Here, the importance of an individual feature is calculated as the relative change in the model’s error when the individual observations of only that feature are shuffled, compared to the error on non-shuffled data. MR thus intentionally breaks the dependence between different correlated features^2^, and is, therefore, a tool to diagnose the model, rather than the data. We use the same loss function as for the model selection, namely ELPD. Since this measure already estimates the logarithm of a quantity of interest (the posterior density), we use the difference between shuffled and non-shuffled ELPD instead of their ratio to estimate the logarithm of the MR for the posterior density. Because we are only interested in qualitatively ranking features by their importance, we normalize the resulting importance value of each feature by their average. To improve the robustness of this measure, the shuffling is repeated multiple times (here, ten times), and the results are averaged. Repeating this procedure for each feature of a model yields positive scores for ranking all features by their importance.

## Results

In this section, we evaluate each of the three models presented above, namely the non-hierarchical model, the model with hierarchical bias term and the full hierarchical model.

### Model predictions

First, we visualize the models’ accuracy and ability to generalize in a model-agnostic way by directly plotting predictions against the corresponding measured “ground-truth” values. For this purpose, we randomly select 80 % of all available data (the training set, shown in blue) to infer model parameters, which we then use to predict the LAI for the remaining 20 % of the data (the test set, shown in green). Due to the probabilistic nature of our models, a full posterior predictive distribution is given for each data point, which we summarize in figure 5 **(A)**,**(C)**, **and** **(E)**. We can observe that all three models make reasonable predictions, i.e., that the predicted LAI grows in proportion to the measured LAI. Because all our generalized linear models assume that the *logarithms* of the LAI scores are homoscedastic, the standard deviation of the predictive distribution increases with the measured LAI, as well. Rather than the raw residuals 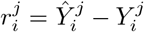, we therefore compute the relative residuals 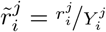,each normalized by the corresponding measured LAI value 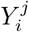, and summarize them in the cumulative histograms shown in figure 5 **(B)**,**(D)** **and (F)**. For all three models, the relative residuals are similar between training and test set, which indicates that they generalize well.

**Figure 5.**
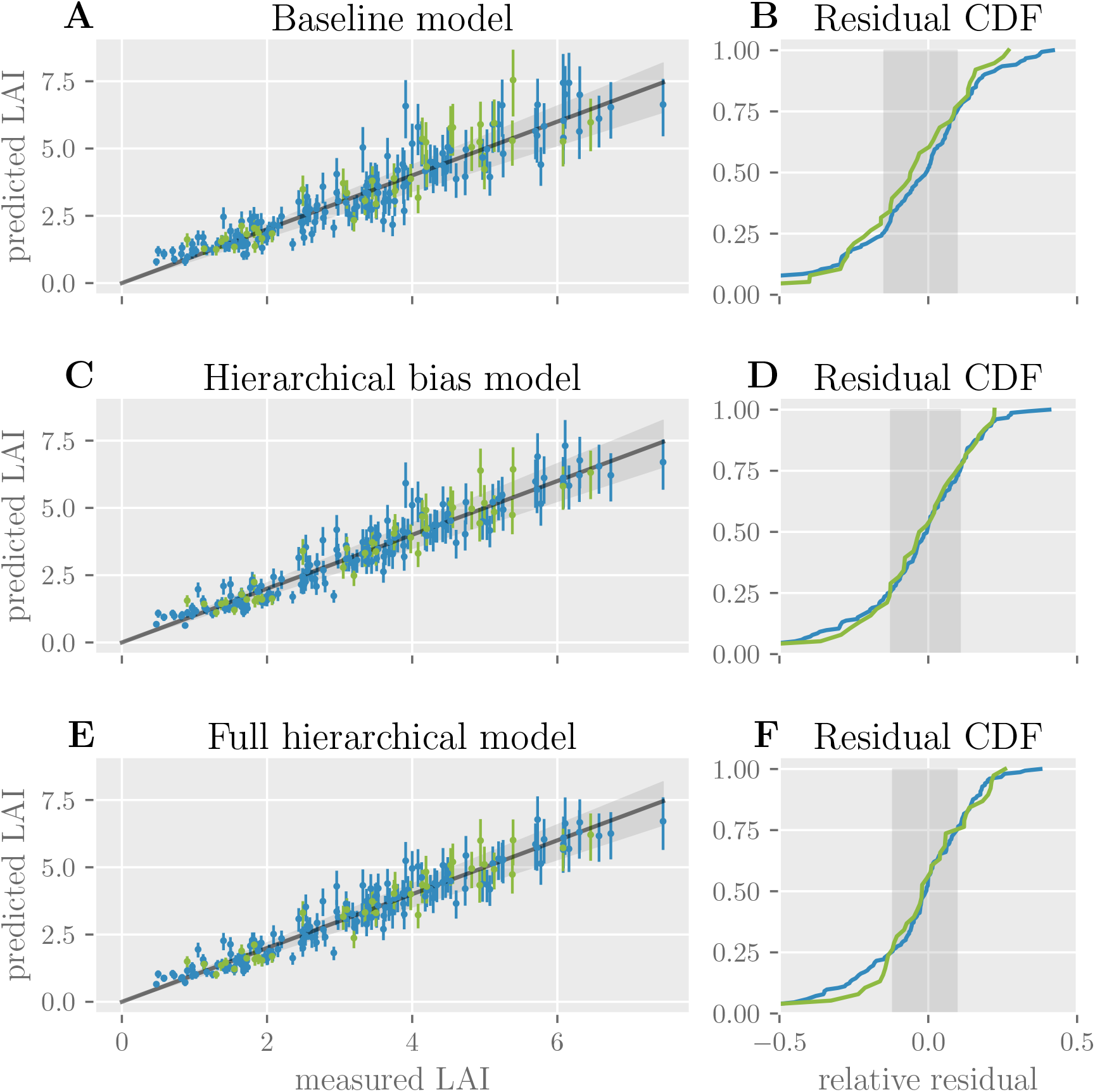
Model predictions of LAI. For the three models, **(A), (C) and (E)** plot for each data point (training data in blue and testing data in green) the predicted LAI values against the actually measured LAI values. Error-bars indicate the interquartile range of the predictive distributions. Dots represent the expected value. The gray line shows the optimal predictions; the best 50% of model predictions lie within the gray cone around it. **(B), (D) and (F)** show the cumulative distribution function of the residuals, each normalized by the corresponding measured LAI value for training and testing data (blue and green lines). The gray areas show the same interquartile range as the cones in **(A), (C) and (E)**.

### Model comparison

To quantify the generalization error of all three models more accurately, we use the PSIS-LOO-CV method on *all* available data to estimate the ELPD on novel data. This procedure yields several highly informative measures, which are summarized in table 2. To verify the convergence of the sampling procedure for each model, we compare the marginal posterior distribution of each parameter across multiple chains and find no discrepancies or divergences (see also S2 Fig, S3 Fig and S4 Fig). We can see that the highest ELPD (indicating the lowest generalization error) is achieved for the two hierarchical models, with little difference between them (− 157.8 and − 157.0, respectively, with a standard deviation of ≈ 11.5 each). Suppose, for the sake of argument, that for a similar dataset we would select models based purely on the ELPD. In that case, it might be a matter of chance whether we would pick the model with only a hierarchical bias term (as in this case) or the full hierarchical model.

**Table 2.** Comparison of the three models using PSIS-LOO-CV. The ELPD ± one standard deviation are listed for each model. *η* denotes the effective number of parameters. For each model, we show the number of data-points for which the Pareto shape parameter *k* falls into either of four different intervals.

|  | Pareto k |  |  |  |  |  |
| --- | --- | --- | --- | --- | --- | --- |
| | ELPD | $\eta$ | (-Inf, 0.5] | (0.5, 0.7] | (0.7, 1] | (1, Inf) |
| model |  |  |  |  |  |  |
| Naive | -185.5±12.2 | 13.3 | 191 | 0 | 0 | 0 |
| Hier. Full | -157.8±11.5 | 24.9 | 180 | 8 | 3 | 0 |
| **Hier. Bias** | **-157.0±11.5** | **15.0** | **187** | **3** | **1** | **0** |

However, due to the limited amount of training data and considering that the number of parameters ranges from 13 for the non-hierarchical baseline model to 17 for the model with hierarchical bias term to 57 for the full hierarchical model, we are also concerned with model complexity and the risk of overfitting. Since LOO-CV estimates generalization error directly (unlike information criteria like AIC or WAIC), it does not need to explicitly penalize a large number of parameters, which is a significant advantage when comparing Bayesian hierarchical models. Instead, it allows us to estimate the model complexity of the three models by the so-called *effective number of parameters η*, which provides some intuition about how many degrees of freedom the model has to approximate the available data. As we see in table 2, *η* = 13.3 is quite close to the parameter count of the non-hierarchical model on the pooled dataset. This only increases slightly to *η* = 15.0 for the hierarchical bias model, despite the fact that it has four additional parameters. However, adding another 44 parameters for the full hierarchical model increases *η* substantially to 24.9.

Since PSIS-LOO-CV emulates conventional LOO-CV, it provides additional information that can help us understand how prone each model is to overfitting: For each data point, the procedure yields the shape parameter *k* of a Pareto distribution, which indicates whether estimating the generalization error for that data point is reliable (*k* ∈ [−∞, 0.7], ideally *k* ≤ 0.5), potentially unreliable (*k* ∈ [0.7, 1]) or entirely unreliable (*k* ∈ [1, ∞]). As table 2 shows, the full hierarchical model struggles with PSIS-LOO-CV for three data points, which may indicate that the model is more prone to overfitting to these potential “outliers” (see also S5 Fig).

As these numbers suggest, the model with a hierarchical bias term is the best choice because it is barely more complex than the non-hierarchical model, yet it performs at least as well as the full hierarchical model.

### An interpretable kernel function

As outlined above, all three models derive their predictions of LAI from a weighted linear combination of features, which we compute by taking inner products between the measured reflectance spectra and a set of B-spline basis functions. These linear operations can be equivalently expressed as taking the inner product between each reflectance spectrum and an inferred *kernel function κ*^*j*^(*λ*), which provides a different, more interpretable perspective on the model.

To motivate this equivalent perspective, we look at how the reflectance spectra affect the linear predictors 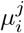 of the respective GLMs in equations 2 to 4, ignoring the contribution of the inferred bias terms here. For all three models^3^, we can use equation 1 to rewrite the contribution of the features extracted from the *i*^th^ reflectance spectrum 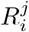 of the *j*^th^ dataset as follows:

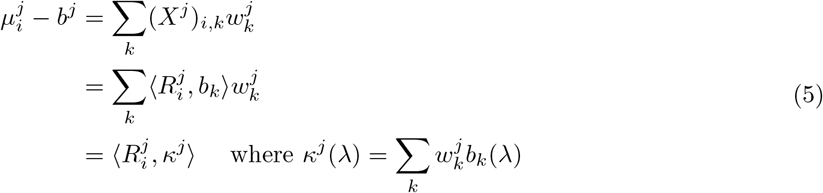

Since the parameters 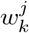 are random variables, the kernel functions *κ*^*j*^ are random variables, samples of which can be generated by combining the (static) basis functions *b*_*k*_ with samples of 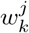. Figure 6 (A) shows the distribution of the inferred kernel function for our model of choice, i.e., the hierarchical bias model (for the other two models, see S6 Fig and S7 Fig). By analyzing this kernel function, we can identify regions of the reflectance spectrum that contribute positively or negatively (e.g., around *λ ≈* 700 nm and *λ ≈* 1300 nm) to the predicted LAI score, and relate them to physical mechanisms.

**Figure 6.**
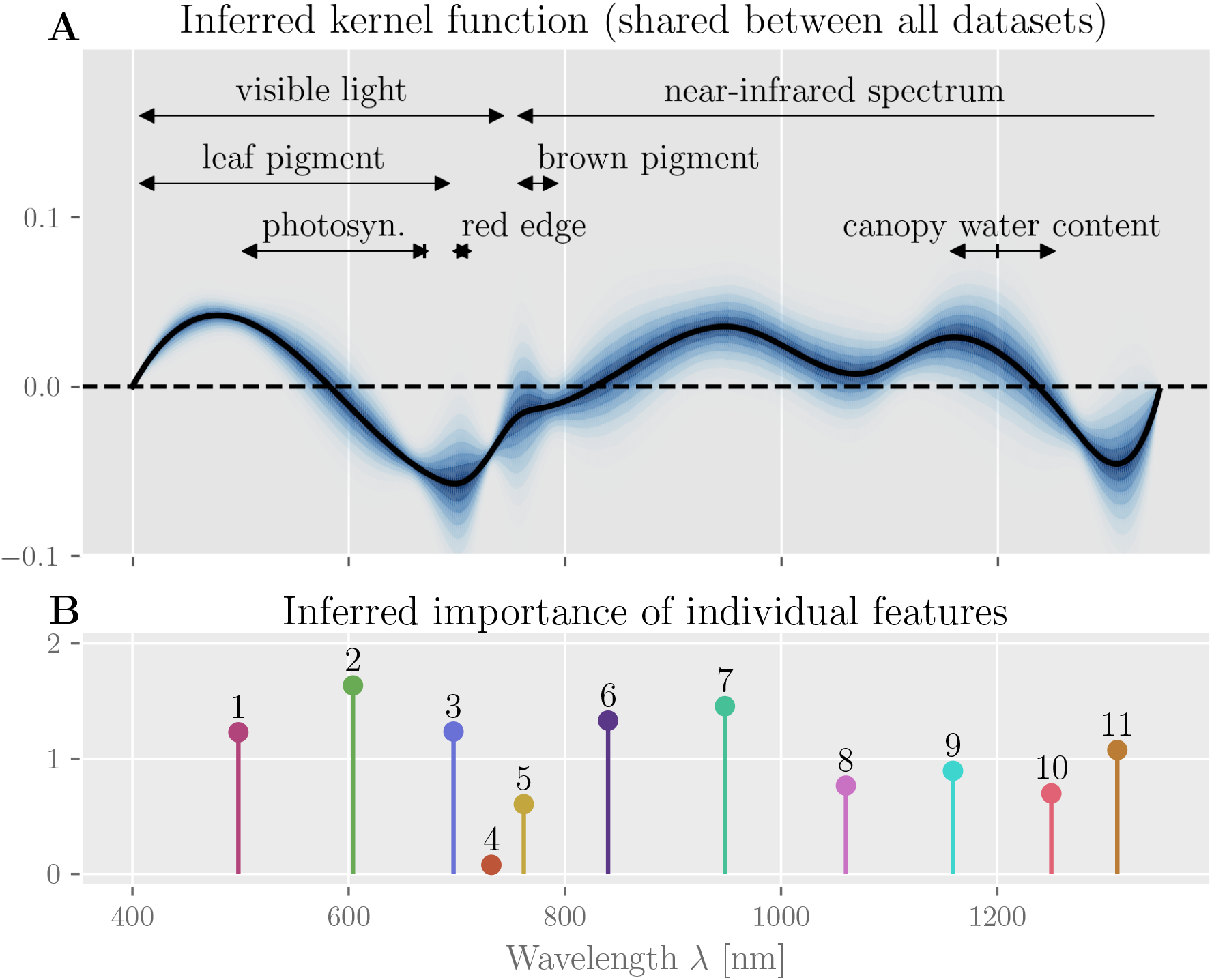
Inferred kernel function and feature importance. **(A)** shows the posterior distribution of the inferred kernel function. The black line represents the expected kernel. We can relate several ranges of the reflectance spectrum to physical phenomena, namely effects due to green leaf pigment (400 nm to 700 nm [42, 43]) and photosynthetic capacity (495 nm to 680 nm, peak at 670 nm [42, 43, 44]) and the red edge region (690 nm to 720 nm [45]) in the visible light range, as well as the canopy’s water content (1150 nm to 1260 nm [46], peak absorption at around 1200 nm [44, 47]) in the near-infrared range. **(B)** shows a stem-plot of the relative importance of each feature (enumerated; normalized by the average feature importance) as well as the resulting estimated importance of each wavelength.

### Feature importance

In addition to the sign and magnitude of each feature’s contribution (which are determined by the inferred weights; c.f. S2 Fig to S4 Fig), we are also interested in how important each individual feature is for the model’s prediction. We quantify this via MR as described above. Figure 6 (B) shows that, with one exception, all features are indeed important for the prediction accuracy of the model. The only notable exception is the fourth feature corresponding to the narrowest basis function *b*_4_ centered around 730 nm (see figure 6 (C)), which indicates that this feature could be removed or an alternative knot-placement procedure could be chosen to reduce model complexity.

## Discussion

In this case study, we investigated how an agronomical variable, the LAI of white winter wheat, can be reliably predicted from measurements of reflectance spectra. The task is challenging because the available dataset is small and consists of multiple heterogeneous sets of measurements carried out at different locations and times. This setting is representative of many problems in data science, particularly in environmental sciences and remote sensing, and Bayesian hierarchical models seem ideally suited to address it [21]. Our results confirm that using a Bayesian hierarchical model not only leads to an improvement in the prediction accuracy over a non-hierarchical approach, but more importantly, it yields several qualitative benefits regarding interpretability, model complexity, and robustness.

One important benefit of the Bayesian hierarchical approach is that an appropriate choice of priors and model structure allows us to integrate additional model parameters without excessively increasing model complexity. For example, the number of spectral features used in our model directly determines the scale of the respective spline basis functions, which determines the resolution of our kernel function. This can create a trade-off between a model with lower spectral resolution and a model with a larger number of parameters. In the Bayesian approach, we can choose the model with more parameters without the risk of overfitting if we formalize our uncertainty and prior assumptions about the parameters appropriately. This is particularly important for hierarchical models, where we might want to add a large number of parameters to account for the specific variations in each subset of the data.

Compare, for example, our full hierarchical model with its 51 parameters to the non-hierarchical baseline model with 13 parameters. Here, the addition of 38 new parameters only increases the effective degrees of freedom of the model by 11 and appears to only moderately increase the risk of overfitting. In our hierarchical bias model, we directly incorporate the fact that each of the four data subsets was recorded at a different growth stage of the plants, which affects the expected LAI, and hence requires a separate bias parameter. However, by simultaneously inferring the shared prior distribution over these separate parameters, we can ensure that the prediction on any one subset of the data benefits from the information contained in all the others. Of course, a non-hierarchical model can also benefit from heterogeneous data (see e.g. [20]), but it may fail in subtle ways if systematic differences *between* the data subsets obscure the relevant associations *within* each dataset ^4^. In general, Bayesian hierarchical models allow us to conveniently include additional information about the dataset, domain knowledge, and regularizing priors, and are particularly useful for small and heterogeneous datasets such as those commonly found in environmental sciences [21].

We employ MCMC sampling to generate unbiased samples of the full posterior distributions over parameters and predictions, which allow us to use additional diagnostic tools and error measures. For example, we can directly estimate posterior densities, credible intervals, and even generalization errors via sample-based methods such as PSIS-LOO-CV, which are more broadly applicable than information criteria such as AIC, WAIC, or BIC [49]. In particular, we saw that a hierarchical model might have a considerably larger number of parameters with a comparatively minor increase in model complexity, making any form of regularization based directly on the number of parameters difficult. Besides better diagnostics, sample-based measures can also provide insights about the data itself, e.g., indicating which data points are potential “outliers” that the model is susceptible to (see S5 Fig).

In addition to descriptive statistics, we also estimate feature importance using an intervention-based model-agnostic method that artificially breaks the dependence between naturally correlated features and thus allows us to infer exactly which features the model relies on for its prediction – independently from the magnitude of the respective parameters. Such information can help domain experts identify potential problems, e.g. if supposedly relevant features are ignored or irrelevant features are relied on. This simple example shows how methods from causal analysis [50] can help explain or interpret the model in qualitatively different ways than descriptive statistics alone.

Because we use a generalized linear model, we can additionally analyze and interpret the model’s linear predictor directly in the measurement space. Since the individual features are extracted from the spectra using B-spline basis functions, this linear predictor is just an inner product between a reflectance spectrum and a kernel function plus an additive bias term. Due to the logarithmic link function, the bias term ultimately has a scaling effect on the LAI predictions. The kernel function directly shows which wavelengths are associated with higher LAI, e.g., short wavelengths of the visible spectrum and much of the near-infrared spectrum, or lower LAI, e.g., around 600 nm to 750 nm. These results can be directly linked to physical phenomena and examined with domain knowledge. For example, the positive association for short wavelengths in the range 400 nm to 550 nm may be attributable to the effect of green leaf pigment, which reflects light in the range 400 nm to 700 nm [42, 43]. Similarly, the pronounced drop around the so-called red-edge region (690 nm to 720 nm [45]), which is related to the plants’ chlorophyll content [51, 52], total nitrogen [53, 54] and yield [55, 56], may be attributable to the plants’ photosynthetic capacity (495 nm to 680 nm [42, 43, 44]) that peaks at around 670 nm.

Finally, we opted for a simplistic, interpretable model of LAI as a function of spectral power, but the hierarchical Bayesian modeling approach makes it easy to extend the proposed model further. One could include more datasets, additional levels of hierarchy (e.g. to extend the model to other related plant species or different geographical regions), or other factors such as soil moisture content [57], the influence of climate change and CO2 concentration on crop growth [15, 16], daily variations in weather due to climate change [58], effects of microclimate [59], the influence of the amount of soil conditioner on the crops [60], and ammonium level in the soil.

## Conclusion

We are convinced that besides accuracy, *interpretability* is a crucial factor for a model to be applied at large, and therefore focus on a simplistic but instructive example here. The choice of priors and hierarchical structure of Bayesian hierarchical models makes it possible to incorporate domain knowledge into the model. Conversely, the linear predictor of our GLM, particularly the inferred kernel, can be analyzed and interpreted by a domain expert. We also use an intervention-based method to estimate feature importance, which can be considered an application of simple causal analysis and provides a qualitatively different perspective that aids interpretability. In addition, we use diagnostic tools such as PSIS-LOO-CV to estimate generalization errors and verify modeling assumptions. Our results show that Bayesian hierarchical models are a powerful and versatile tool for environmental sciences, especially when the available data is scarce and comes from heterogeneous sources.

## Author Contributions

**OS and JL** Conceptualization, Methodology, Project administration, Software, Visualization, Writing – original draft

**GP** Supervision, Resources, Conceptualization, Validation, Writing – review & editing

**BS and TJ** Data curation, Conceptualization, Validation, Writing – review & editing

## Supporting Information

**S1 Fig.**
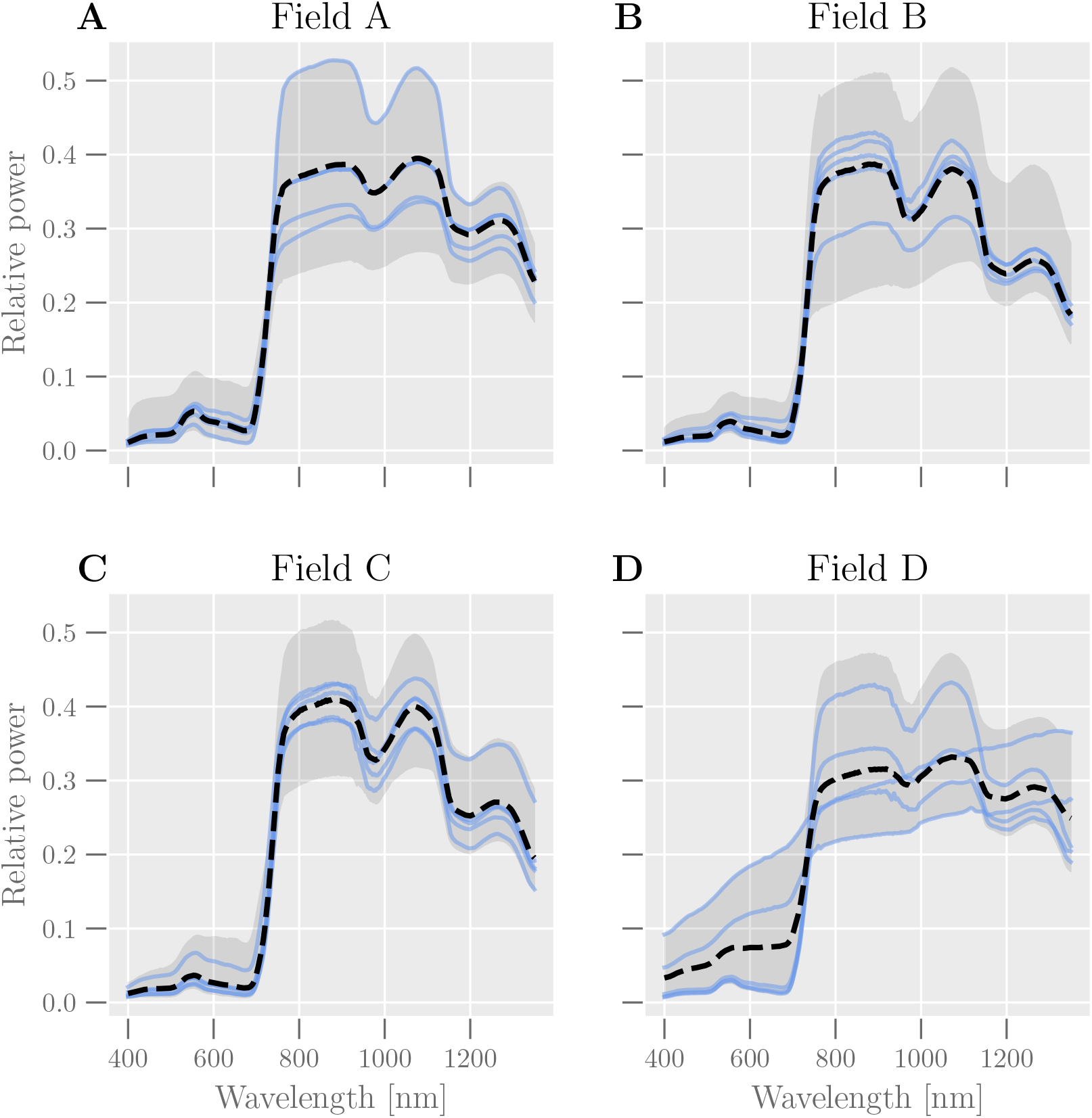
Reflectance spectra from the four datasets. Solid lines show five randomly selected spectra from each dataset. The dashed lines show the average for the respective dataset, and the gray regions show the value range (from minimum to maximum) for each wavelength.

**S2 Fig.**
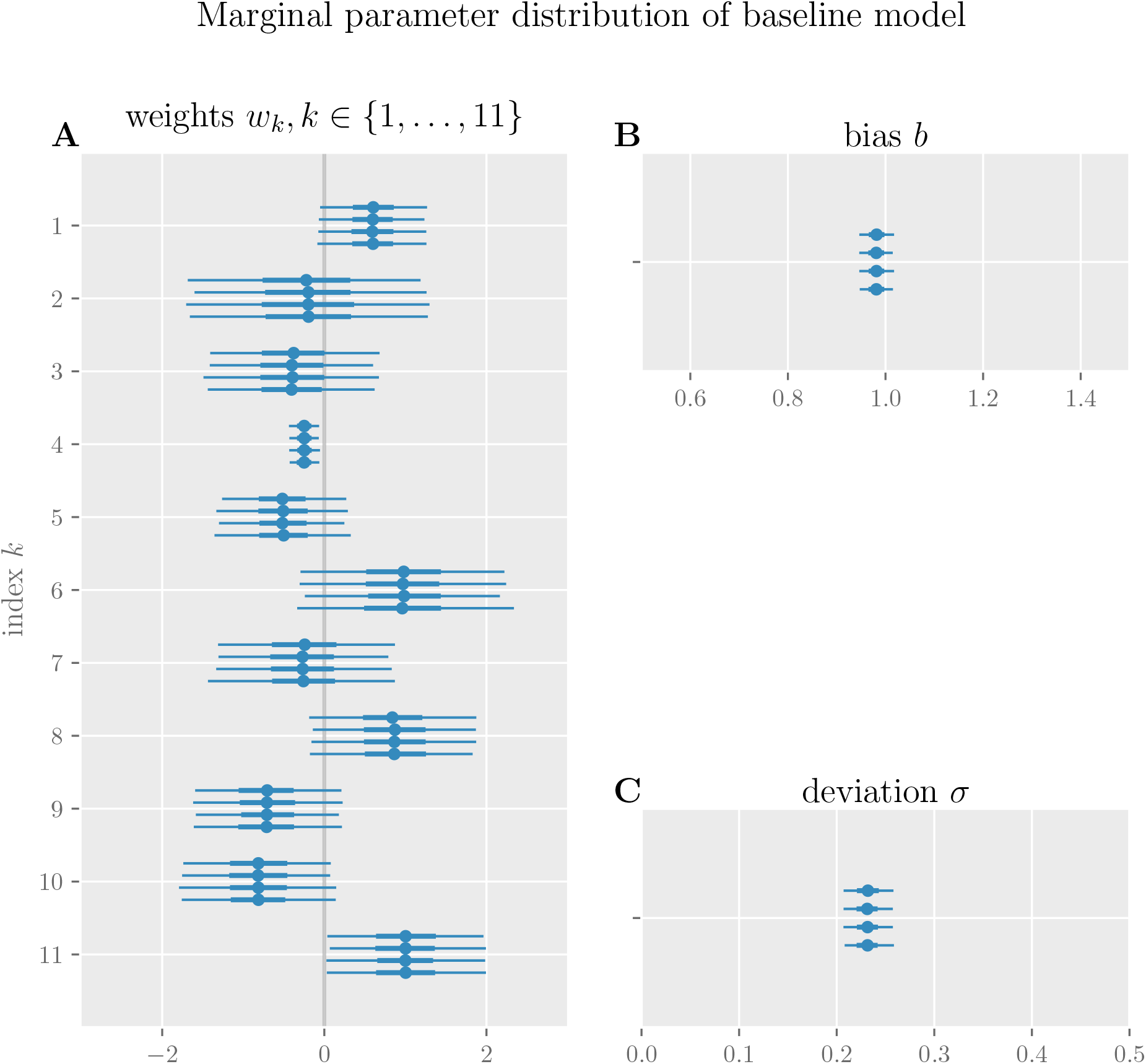
Marginal posterior distributions of all parameters of the baseline model. For each of four Markov chains, the mean (dot), the interquartile range from the 25 % to 75 % quantile (thick horizontal lines) as well as the 2.5 % to 97.5 % quantile (thin horizontal lines) are shown. For all parameters, these summary statistics of the marginal distribution are similar across all four chains, indicating convergence of the MCMC sampling scheme.

**S3 Fig.**
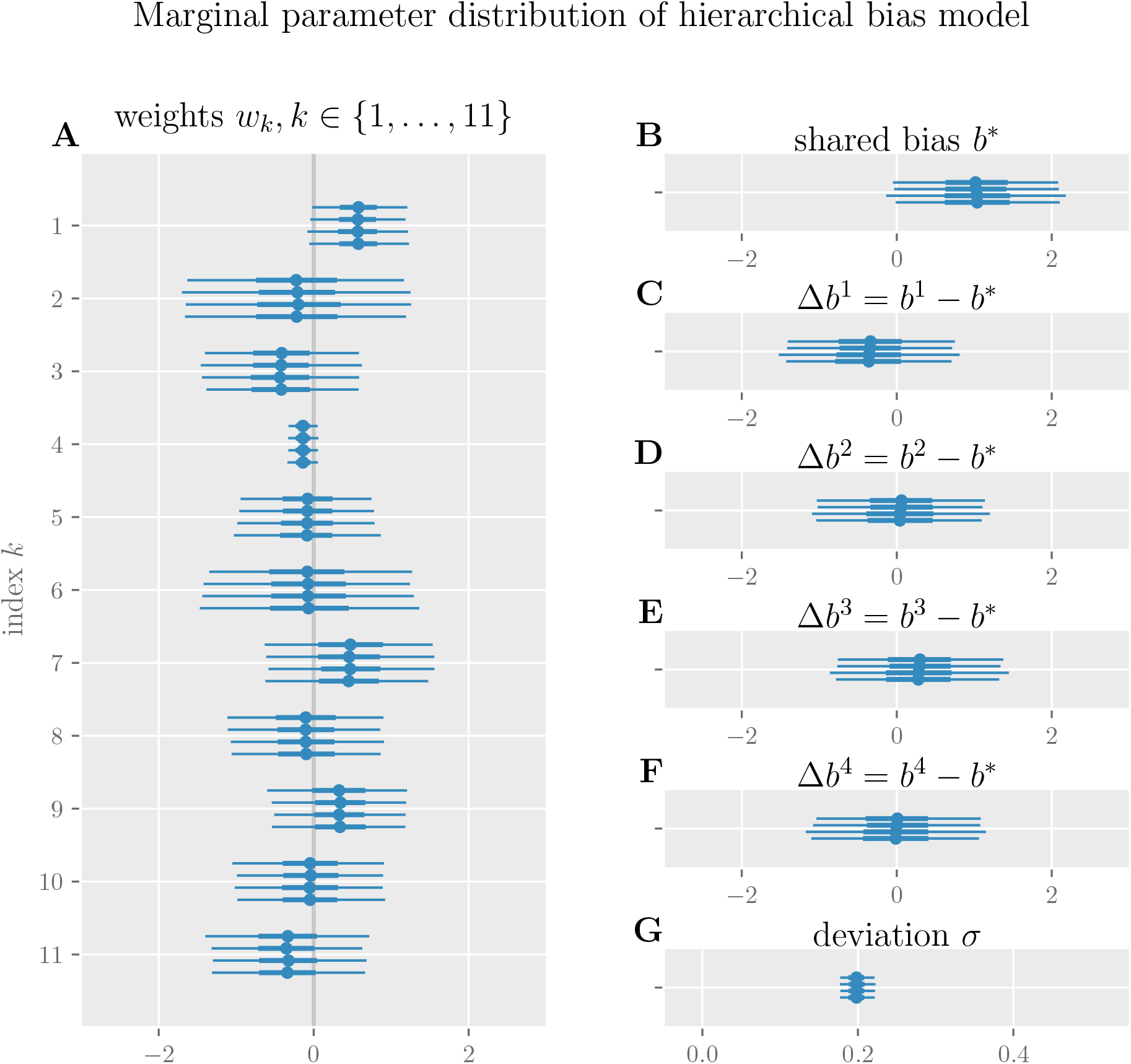
Marginal posterior distributions of all parameters of the hierarchical bias model. For each of four Markov chains, the mean (dot), the interquartile range from the 25 % to 75 % quantile (thick horizontal lines) as well as the 2.5 % to 97.5 % quantile (thin horizontal lines) are shown. (C) to (F) show the differences between the shared bias parameter *b*^*^ and the dataset-specific bias parameters *b*^*j*^. For all parameters, these summary statistics of the marginal distribution are similar across all four chains, indicating convergence of the MCMC sampling scheme.

**S4 Fig.**
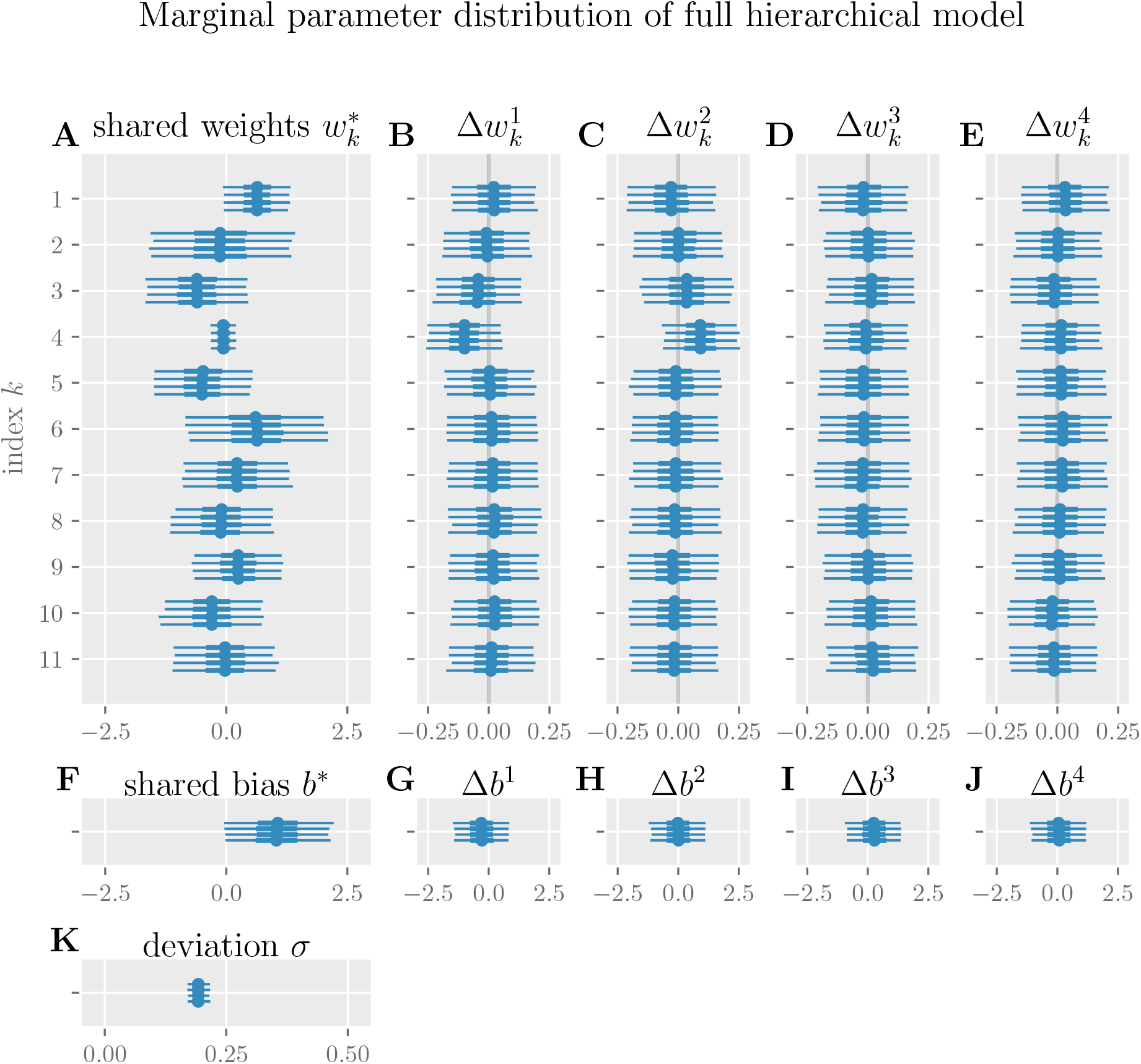
Marginal posterior distributions of all parameters of the full hierarchical model. For each of four Markov chains, the mean (dot), the interquartile range from the 25 % to 75 % quantile (thick horizontal lines) as well as the 2.5 % to 97.5 % quantile (thin horizontal lines) are shown. (B) to (E) show the differences between the shared weight parameters 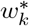 and the dataset-specific weight parameters 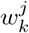, and (G) to (J) show the differences between the shared bias parameter *b*^*^ and the dataset-specific bias parameters *b*^*j*^. For all parameters, these summary statistics of the marginal distribution are similar across all four chains, indicating convergence of the MCMC sampling scheme.

**S5 Fig.**
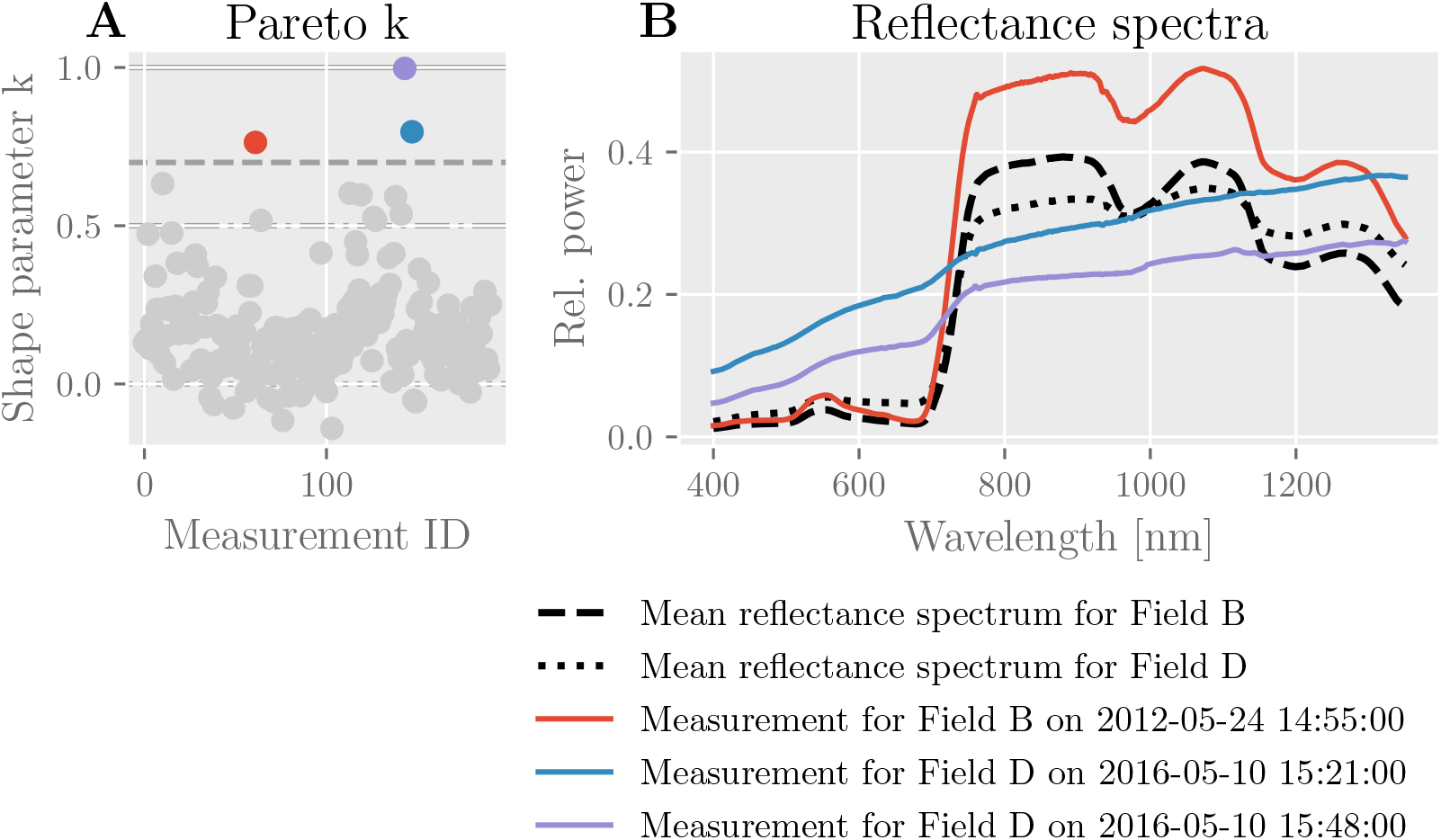
Evaluation of PSIS-LOO-CV diagnostic for full hierarchical model. (A) shows the shape parameters (Pareto k) computed for each datapoint by the PSIS-LOO-CV method for the full hierarchical model. For three measurements (color-coded), the shape parameter exceeds the critical value of 0.7 and PSIS-LOO-CV becomes unreliable. (B) shows the reflectance spectra corresponding to these three datapoints (solid lines, color-coded) as well as the mean reflectance spectra of the respective datasets (dashed lines).

**S6 Fig.**
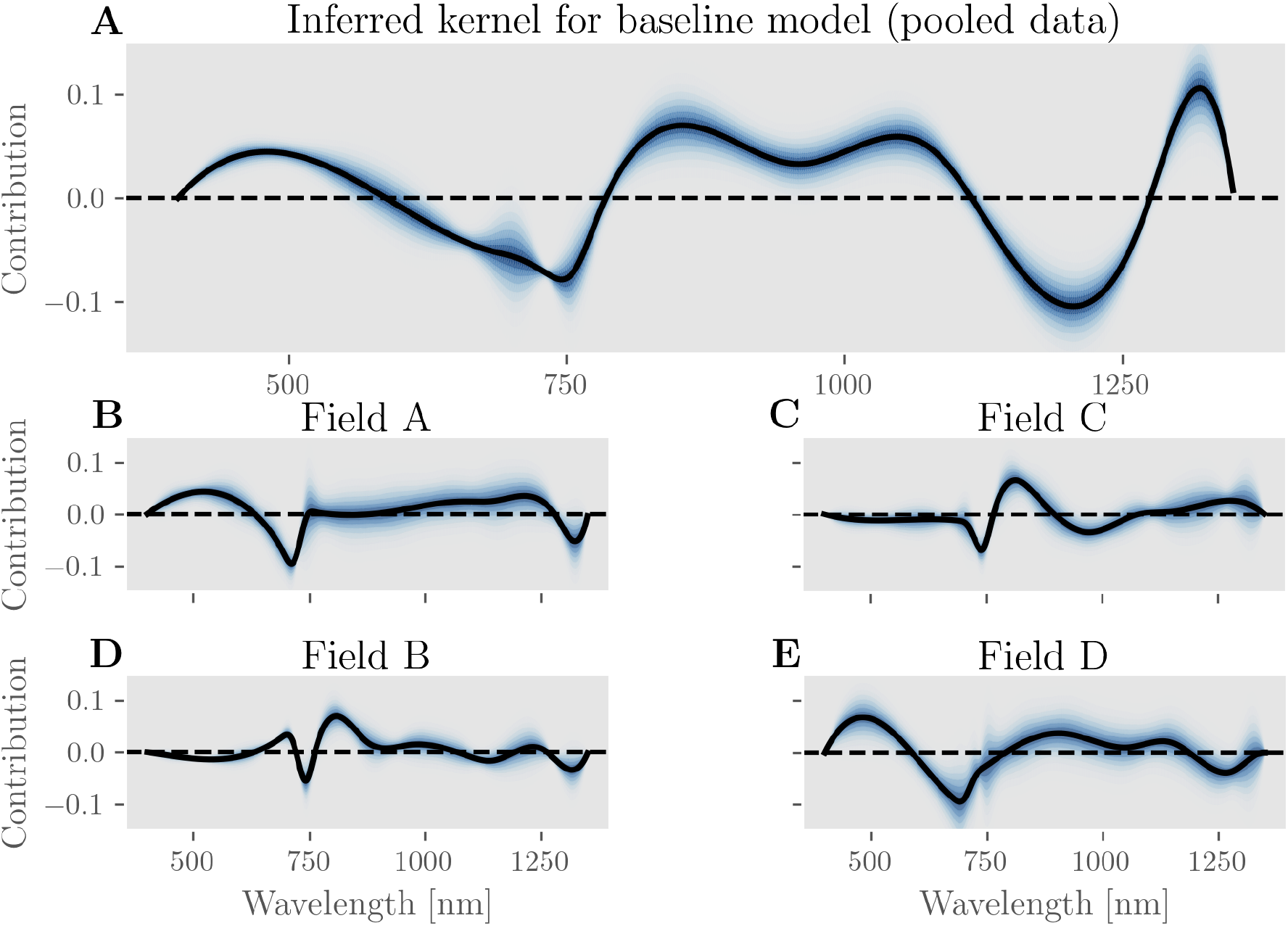
Inferred kernel for baseline model. (A) shows the kernel of the baseline model when fit to the entire pooled dataset. For reference, (B) through (E) show the different kernels that the baseline model would infer from each of the four datasets in isolation. We can see large, qualitative differences between the five shown kernel functions. In particular, the pooled data results in a kernel function with a sizeable dip around a wavelength of 1200 nm, which is entirely absent from any of the kernels inferred for the individual datasets. This indicates that systematic differences between the datasets might introduce spurious associations between spectral features and LAI predictions, which pose a risk for misinterpretation. This effect is avoided entirely by a full hierarchical model (see S7 Fig) and much reduced by the hierarchical bias model (see figure 6).

**S7 Fig.**
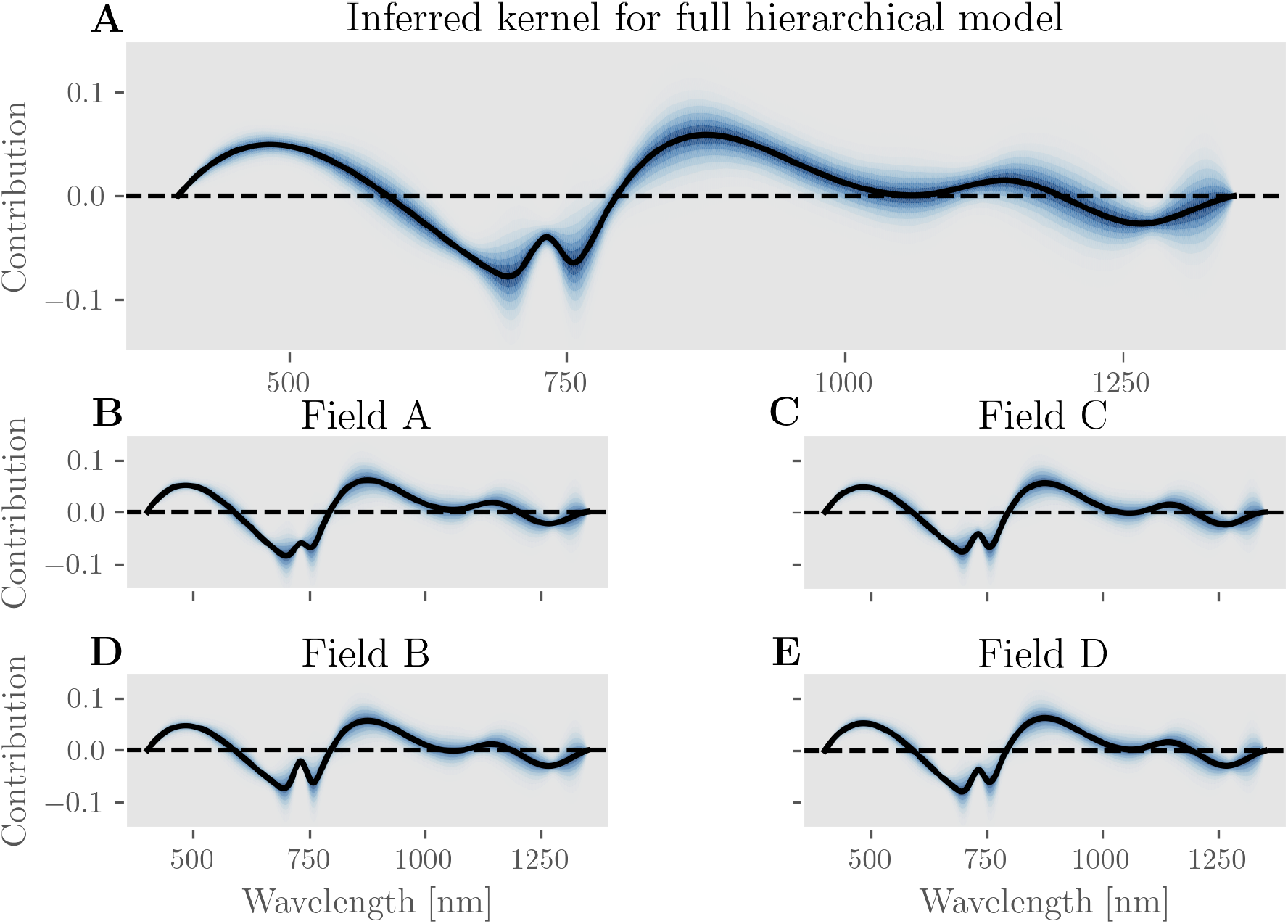
Inferred kernel for full hierarchical model. (A) shows the shared kernel function of the full hierarchical model, and (B) through (E) show the specific kernel functions inferred for each dataset. The inferred dataset-specific kernels deviate only little from the shared kernel, yet in contrast to the baseline model in S6 Fig, there is no pronounced dip around the wavelength 1200 nm.

## Footnotes

1 This is equivalent to computing the curvature of the smoothed average reflectance spectrum 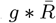.

2 In our model, different features are computed by taking the inner product between the reflectance spectra and a set of overlapping not independent basis functions, and are hence certainly correlated.

3 To simplify notation, we write *w*^*j*^ and *b*^*j*^ for the (possibly) dataset specific weight and bias terms, and set *w*^*j*^ = *w* or *b*^*j*^ = *b* for models that don’t make these dataset specific distinctions.

4 In S6 Fig we show that this is indeed the case here, as wavelength around 1200nm lead to a pronounced dip in the kernel function when data from multiple datasets is pooled, but this association disappears if the model is instead fit to any individual dataset. Looking at the pooled dataset, we would therefore be led to conclude that lower spectral power around 1200nm is a strong predictor of higher LAI. While this is correct on the artificially pooled dataset, it appears to be incorrect on any of the individual datasets. This is an instance of Simpson’s paradox [48], which suggests that a hierarchical model is more appropriate here

## Notes

https://github.com/ostojanovic/bayesian_lai

### Competing Interest Statement

The authors have declared no competing interest.

## References

1. D. J. Watson. Comparative Physiological Studies on the Growth of Field Crops: I. Variation in Net Assimilation Rate and Leaf Area between Species and Varieties, and within and between Years. Annals of Botany, 11(1):41–76, January 1947.

2. J. M. Chen and T. A. Black. Defining leaf area index for non-flat leaves. Plant, Cell & Environment, 15(4):421–429, 1992.

3. J. Montheith and M. Unsworth. Principles of Environmental Physics. Academic Press, San Diego, CA, USA, 2007.

4. Nathalie J.J. Bréda. Ground-based measurements of leaf area index: a review of methods, instruments and current controversies. Journal of Experimental Botany, 54(392):2403–2417, November 2003.

5. Guangjian Yan, Ronghai Hu, Jinghui Luo, Marie Weiss, Hailan Jiang, Xihan Mu, Donghui Xie, and Wuming Zhang. Review of indirect optical measurements of leaf area index: Recent advances, challenges, and perspectives. Agricultural and Forest Meteorology, 265:390–411, February 2019.

6. Sidney Cox. Information technology: The global key to precision agriculture and sustainability. Computers and Electronics in Agriculture, 36:93–111, 11 2002.

7. David J. Mulla. Twenty five years of remote sensing in precision agriculture: Key advances and remaining knowledge gaps. Biosystems Engineering, 114(4):358 – 371, 2013. Special Issue: Sensing Technologies for Sustainable Agriculture.

8. John K. Schueller. A review and integrating analysis of Spatially-Variable Control of crop production. Fertilizer research, 33(1):1–34, October 1992.

9. Laurent Kergoat, Sébastien Lafont, Hervé Douville, Béatrice Berthelot, Gérard Dedieu, Serge Planton, and Jean-François Royer. Impact of doubled co2 on global-scale leaf area index and evapotranspiration: Conflicting stomatal conductance and lai responses. Journal of Geophysical Research: Atmospheres, 107(D24):ACL 30–1–ACL 30–16, 2002.

10. H. Yan, S.Q. Wang, D. Billesbach, W. Oechel, J.H. Zhang, T. Meyers, T.A. Martin, R. Matamala, D. Baldocchi, G. Bohrer, D. Dragoni, and R. Scott. Global estimation of evapotranspiration using a leaf area index-based surface energy and water balance model. Remote Sensing of Environment, 124:581 – 595, 2012.

11. Simic Milas Anita, Fernandes Richard, and Shusen Wang. Assessing the impact of leaf area index on evapotranspiration and groundwater recharge across a shallow water region for diverse land cover and soil properties. Journal of Water Resource and Hydraulic Engineering, 3:60–73, 12 2014.

12. Gregory P. Asner, Jonathan M. O. Scurlock, and Jeffrey A. Hicke. Global synthesis of leaf area index observations: implications for ecological and remote sensing studies. Global Ecology and Biogeography, 12(3):191–205, 2003.

13. N.H Broge and J.V Mortensen. Deriving green crop area index and canopy chlorophyll density of winter wheat from spectral reflectance data. Remote Sensing of Environment, 81(1):45–57, 2002.

14. M. S. Moran, S. J. Maas, and P. J. Pinter Jr. Combining remote sensing and modeling for estimating surface evaporation and biomass production. Remote Sensing Reviews, 12(3-4):335–353, 1995.

15. Zhiqiang Gao, Wei Gao, and James Slusser. The response of leaf area index to climate change during 1981-2000 in China. In Remote Sensing and Modeling of Ecosystems for Sustainability II, volume 5884, page 58840S. International Society for Optics and Photonics, September 2005.

16. Anthony Manea and Michelle R. Leishman. Leaf Area Index Drives Soil Water Availability and Extreme Drought-Related Mortality under Elevated CO2 in a Temperate Grassland Model System. PLoS ONE, 9(3), March 2014.

17. Inge Jonckheere, Stefan Fleck, Kris Nackaerts, Bart Muys, Pol Coppin, Marie Weiss, and Frédéric Baret. Review of methods for in situ leaf area index determination: Part i. theories, sensors and hemispherical photography. Agricultural and Forest Meteorology, 121(1):19–35, 2004.

18. Juanjuan Zhang, Tao Cheng, Wei Guo, Xin Xu, Hongbo Qiao, Yimin Xie, and Xinming Ma. Leaf area index estimation model for UAV image hyperspectral data based on wavelength variable selection and machine learning methods. Plant Methods, 17(1):49, 2021.

19. K.H. Kjaer, K.K. Petersen, and M. Bertelsen. Protective rain shields alter leaf microclimate and photosynthesis in organic apple production. Acta Horticulturae, 1134:317–326, 2016.

20. Bastian Siegmann and Thomas Jarmer. Comparison of different regression models and validation techniques for the assessment of wheat leaf area index from hyperspectral data. International Journal of Remote Sensing, 36(18):4519–4534, 2015.

21. Gregory L. Britten, Yara Mohajerani, Louis Primeau, Murat Aydin, Catherine Garcia, Wei-Lei Wang, Benoîît Pasquier, B. B. Cael, and François W. Primeau. Evaluating the benefits of bayesian hierarchical methods for analyzing heterogeneous environmental datasets: A case study of marine organic carbon fluxes. Frontiers in Environmental Science, 9:28, 2021.

22. Cornelius Senf, Dirk Pflugmacher, Marco Heurich, and Tobias Krueger. A bayesian hierarchical model for estimating spatial and temporal variation in vegetation phenology from landsat time series. Remote Sensing of Environment, 194:155–160, 2017.

23. Xiaojie Gao, Josh M. Gray, and Brian J. Reich. Long-term, medium spatial resolution annual land surface phenology with a bayesian hierarchical model. Remote Sensing of Environment, 261:112484, 2021.

24. Chad Babcock, Andrew O. Finley, and Nathaniel Looker. A bayesian model to estimate land surface phenology parameters with harmonized landsat 8 and sentinel-2 images. Remote Sensing of Environment, 261:112471, 2021.

25. Adam M. Wilson, John A. Silander, Alan Gelfand, and Jonathan H. Glenn. Scaling up: linking field data and remote sensing with a hierarchical model. International Journal of Geographical Information Science, 25(3):509–521, 2011.

26. Luqi Xing, Xuejian Li, Huaqiang Du, Guomo Zhou, Fangjie Mao, Tengyan Liu, Junlong Zheng, Luofan Dong, Meng Zhang, Ning Han, Xiaojun Xu, Weiliang Fan, and Di’en Zhu. Assimilating multiresolution leaf area index of moso bamboo forest from modis time series data based on a hierarchical bayesian network algorithm. Remote Sensing, 11(1), 2019.

27. Kusum J. Naithani, Doug C. Baldwin, Katie P. Gaines, Henry Lin, and David M. Eissenstat. Spatial distribution of tree species governs the spatio-temporal interaction of leaf area index and soil moisture across a forested landscape. PLOS ONE, 8(3):1–12, 03 2013.

28. Daniel Schraik, Petri Varvia, Lauri Korhonen, and Miina Rautiainen. Bayesian inversion of a forest reflectance model using sentinel-2 and landsat 8 satellite images. Journal of Quantitative Spectroscopy and Radiative Transfer, 233:1–12, 2019.

29. X.Q. Xu, J.S. Lu, N. Zhang, T.C. Yang, J.Y. He, X. Yao, T. Cheng, Y. Zhu, W.X. Cao, and Y.C. Tian. Inversion of rice canopy chlorophyll content and leaf area index based on coupling of radiative transfer and bayesian network models. ISPRS Journal of Photogrammetry and Remote Sensing, 150:185–196, 2019.

30. Rogério P. Soratto, Aline O. Matoso, Amanda P. Gilabel, Fabiana M. Fernandes, Rai A. Schwalbert, and Ignacio A. Ciampitti. Agronomic optimal plant density for semiupright cowpea as a second crop in southeastern brazil. Crop Science, 60(5):2695–2708, 2020.

31. Yonghua Qu, Jindi Wang, Huawei Wan, Xiaowen Li, and Guoqing Zhou. A bayesian network algorithm for retrieving the characterization of land surface vegetation. Remote Sensing of Environment, 112(3):613–622, 2008.

32. Carl de Boor. A Practical Guide to Splines. Applied Mathematical Sciences. Springer-Verlag, New York, 1978.

33. Gordon Pipa, Zhe Chen, Sergio Neuenschwander, Bruss Lima, and Emery N. Brown. Mapping of Visual Receptive Fields by Tomographic Reconstruction. Neural computation, 24(10):2543–2578, October 2012.

34. Tegoeh Tjahjowidodo, VT Dung, and ML Han. A fast non-uniform knots placement method for b-spline fitting. In 2015 IEEE International Conference on Advanced Intelligent Mechatronics (AIM), pages 1490–1495. IEEE, 2015.

35. Lluís Jordi Ferrer Arnau, Ramón Reig-Bolaño, Pere Martí-Puig, Amalia Manjabacas, and Vicenç Parisi-Baradad. Efficient cubic spline interpolation implemented with fir filters. International Journal of Computer Information Systems and Industrial Management Applications, 105:98–105, 2013.

36. Andrew Gelman, John B Carlin, Hal S Stern, David B Dunson, Aki Vehtari, and Donald B Rubin. Bayesian data analysis. CRC press, 2013.

37. P McCullagh and John A Nelder. Generalized Linear Models, volume 37. CRC Press, 1989.

38. Matthew D. Hoffman and Andrew Gelman. The No-U-Turn Sampler: Adaptively Setting Path Lengths in Hamiltonian Monte Carlo. arXiv:1111.4246 [cs, stat], November 2011. arXiv: 1111.4246.

39. John Salvatier, Thomas V. Wiecki, and Christopher Fonnesbeck. Probabilistic programming in Python using PyMC3. PeerJ Computer Science, 2:e55, April 2016.

40. Aki Vehtari, Daniel Simpson, Andrew Gelman, Yuling Yao, and Jonah Gabry. Pareto smoothed importance sampling. arXiv:1507.02646 [stat], 2019.

41. Aaron Fisher, Cynthia Rudin, and Francesca Dominici. All Models are Wrong, but Many are Useful: Learning a Variable’s Importance by Studying an Entire Class of Prediction Models Simultaneously. arXiv:1801.01489 [stat], 2019. arXiv: 801.01489 version: 5.

42. Yan-Ping Cen and Janet F. Borman. The Response of Bean Plants to UV-B Radiation Under Different Irradiances of Background Visible Light. Journal of Experimental Botany, 41(11):1489–1495, 11 1990.

43. Nieves Aparicio, Dolors Villegas, Jaume Casadesus, José Luis Araus, and Conxita Royo. Spectral vegetation indices as nondestructive tools for determining durum wheat yield. Agronomy Journal, 92(1):83–91, 2000.

44. Vivian Roca Schwendler Weber, José Luis Araus, Jill E. Cairns, Ciro Dagnny Arce Séanchez, Albrecht E Melchinger, and Elena Orsini. Prediction of grain yield using reflectance spectra of canopy and leaves in maize plants grown under different water regimes. Field Crops Research, 128, 2012.

45. Jianfeng Zhang, Wenting Han, Lvwen Huang, Zhiyong Zhang, Yimian Ma, and Yamin Hu. Leaf chlorophyll content estimation of winter wheat based on visible and near-infrared sensors. Sensors (Basel, Switzerland), 16, 03 2016.

46. Daniel Sims and John Gamon. Estimation of vegetation water content and photosynthetic tissue area from spectral reflectance: A comparison of indices based on liquid water and chlorophyll absorption features. Remote Sensing of Environment, 84:526–537, 04 2003.

47. Chao Wang, Meichen Feng, Wude Yang, Guangwei Ding, Lujie Xiao, Guangxin Li, and Tingting Liu. Extraction of sensitive bands for monitoring the winter wheat (triticum aestivum) growth status and yields based on the spectral reflectance. PLOS ONE, 12(1):1–16, 01 2017.

48. Judea Pearl. Comment: understanding simpson’s paradox. The American Statistician, 68(1):8–13, 2014.

49. Aki Vehtari, Andrew Gelman, and Jonah Gabry. Practical Bayesian model evaluation using leave-one-out cross-validation and WAIC. Statistics and Computing, 27(5):1413–1432, September 2017. arXiv: 1507.04544.

50. Judea Pearl et al. Causal inference in statistics: An overview. Statistics surveys, 3:96–146, 2009.

51. Andrew Richardson, Shane P. Duigan, and Graeme Berlyn. An evaluation of noninvasive methods to estimate foliar chlorophyll content. New Phytologist, 153:185 – 194, 01 2002.

52. Liangyun Liu, Jihua Wang, Wenjiang Huang, Chunjiang Zhao, Bing Zhang, and Qingxi Tong. Estimating winter wheat plant water content using red edge parameters. International Journal of Remote Sensing, 25(17):3331–3342, 2004.

53. J.G.P.W. Clevers and Lammert Kooistra. Using hyperspectral remote sensing data for retrieving canopy water content. WHISPERS ’09 - 1st Workshop on Hyperspectral Image and Signal Processing: Evolution in Remote Sensing, pages 1–4, 09 2009.

54. Yisong Cheng, Chunsheng Hu, Hui Dai, and Yuping Lei. Spectral red edge parameters for winter wheat under different nitrogen support levels. Proceedings of SPIE - The International Society for Optical Engineering, 08 2005.

55. Vali Rasooli Sharabiani, Noboru Noguchi, and Kazunobu Ishi. Significant wavelengths for prediction of winter wheat growth status and grain yield using multivariate analysis. Engineering in Agriculture, Environment and Food, 7:14–21, 02 2014.

56. Wenjiang Huang, Jindi Wang, Huawei Wan, Jihua Wang, Liangyun Liu, and Chunjiang Zhao. Application of red edge variables in winter wheat nutrition diagnosis. In IGARSS 2004. 2004 IEEE International Geoscience and Remote Sensing Symposium, volume 6, pages 4052–4055 vol.6, 2004.

57. M.H. Rad, M.H. Assare, Mohammad Banakar, and Maziar Soltani. Effects of Different Soil Moisture Regimes on Leaf Area Index, Specific Leaf Area and Water use Efficiency in Eucalyptus (Eucalyptus camaldulensis Dehnh) under Dry Climatic Conditions. Asian Journal of Plant Sciences, 10:294–300, 2011.

58. Daniel T. C. Cox, Ilya M. D. Maclean, Alexandra S. Gardner, and Kevin J. Gaston. Global variation in diurnal asymmetry in temperature, cloud cover, specific humidity and precipitation and its association with leaf area index. Global Change Biology, n/a(n/a), 2020.

59. Stephen R. Hardwick, Ralf Toumi, Marion Pfeifer, Edgar C. Turner, Reuben Nilus, and Robert M. Ewers. The relationship between leaf area index and microclimate in tropical forest and oil palm plantation: Forest disturbance drives changes in microclimate. Agricultural and Forest Meteorology, 201:187–195, February 2015.

60. Lijun Su, Quanjiu Wang, Chunxia Wang, and Yuyang Shan. Simulation Models of Leaf Area Index and Yield for Cotton Grown with Different Soil Conditioners. PLoS ONE, 10(11), November 2015.

